# ceTDP: A Practical Tutorial on Valid and Quantitative Cluster-Extent Inference in Neuroimaging

**DOI:** 10.64898/2026.11.15.766765

**Authors:** Xu Chen, Lucas Peek, Ramin Monajemi, Paweł Górecki, Thomas E. Nichols, Jelle J. Goeman, Wouter D. Weeda

## Abstract

We introduce ceTDP, a two-stage framework for cluster-extent inference in neuroimaging that extends conventional random field theory-based and permutation-based methods. In contrast to standard cluster inference, which yields binary significance, ceTDP additionally quantifies the proportion of truly active voxels within clusters using the true discovery proportion (TDP). The method is built on closed testing and provides strong family-wise error rate control while remaining consistent with standard cluster inference.

The framework operates in two stages: estimation of a cluster-extent threshold, followed by computation of the TDP lower bound. Two complementary procedures are provided: a conservative, statistically guaranteed TDP lower bound (ceTDP-cons) and a heuristic approach for obtaining a potentially tighter estimate of the TDP lower bound (ceTDP-heur). Together, they enable both rigorous inference and improved sensitivity in cluster-level analysis.

## 1 Introduction

Statistical inference in neuroimaging is challenged by the multiple comparisons problem, as most studies typically involve testing hundreds of thousands of voxels. The most commonly used parametric cluster-extent inference, which is based on Gaussian random field theory (RFT), has become standard by testing the significance of clusters of contiguous voxels instead of individual voxels to offer greater sensitivity. As implemented in the widely used neuroimaging software tools, such as Statistical Parametric Mapping (SPM; Worsley et al., 1992; Friston et al., 1994; Worsley et al., 1996) and FMRIB

Software Library (FSL; Smith et al., 2004; Jenkinson et al., 2012), RFT-based cluster-extent inference provides a practical solution for controlling the family-wise error rate (FWER) at the cluster level. In addition to these parametric methods, permutation-based nonparametric cluster inference (Hayasaka and Nichols, 2003; Hayasaka et al., 2004) serves as an alternative that does not rely on the RFT-related distributional assumptions, and provides more robust and reliable inference when these assumptions are violated, especially when sample size is small.

Although the established cluster extent inference framework remains practically useful, several studies have revealed critical statistical flaws (Woo et al., 2014; Eklund et al., 2016; Rosenblatt et al., 2018; Botvinik-Nezer et al., 2020). A crucial limitation is the spatial specificity paradox: detecting a significant cluster only allows the conclusion that at least one voxel within it is active (Woo et al., 2014; Rosenblatt et al., 2018). This leads to the common misinterpretation that the statistical significance of a cluster implies that all voxels within that cluster are significantly activated (Woo et al., 2014; Rosenblatt et al., 2018). Such misunderstanding limits the ability to make reliable inferences about activation localization.

To address the issues with spatial specificity, we introduce the ceTDP methodology (Goeman et al., 2023), which is a robust extension of traditional RFT-based or permutation-based cluster-extent inference that incorporates a closed testing framework. This approach provides strong cluster-level FWER control while simultaneously calculating lower confidence bounds for the true discovery proportion (TDP), i.e., the proportion of truly activated voxels within a given cluster. This advances statistical inference from a common binary decision based on a *p*-value to a quantitative assessment of true activation proportion using the TDP.

The TDP is defined as the proportion of truly activated voxels within a region, which may be anatomically predefined, user-specified, or data-driven (e.g., supra-threshold clusters, STCs). It provides a quantitative and interpretable measure of the activation strength in a region. By constructing simultaneous lower confidence bounds for the TDP that hold for all possible regions, one can control the FWER at cluster level across any set of regions. Specifically, ensuring that the TDP of any claimed region is greater than zero is statistically equivalent to controlling cluster-level FWER. This tutorial revisits the theoretical background and introduces practical implementation of the ceTDP framework, with the aim of making this inference approach more accessible to the neuroimaging community.

The rest of this paper is organized as follows. Section 2 provides an overview of the ceTDP methodology, followed by a motivating example in Section 3. Section 4 introduces the two core stages of the ceTDP framework. Building upon this, Section 5 provides a step-by-step workflow for implementing the entire analysis. Section 6 outlines best practices and common pitfalls to ensure valid analysis. Finally, we conclude with a discussion of its impact in Section 7.

## 2 Overview of ceTDP Methodology

This section provides a non-technical overview of the ceTDP methodology. For a more thorough theoretical introduction, see Goeman et al. (2023). We begin by outlining the main limitations of conventional RFT-based inference, then introduce the closed testing solution, highlight the benefits of quantifying the TDP, and conclude with an efficient two-stage framework that makes the approach practical.

The ceTDP framework offers two complementary procedures for quantifying the TDP lower bound: ceTDP-cons, which provides a statistically guaranteed conservative lower bound, and ceTDP-heur, which uses heuristic optimization to obtain an estimate of the TDP lower bound.

### 2.1 Flaws in Standard Cluster Inference

Standard RFT-based cluster-extent inference, as implemented in SPM and FSL, suffers from well-documented statistical flaws, including inflated false positive rates, poor spatial specificity, and circular inference (Woo et al., 2014; Eklund et al., 2016; Rosenblatt et al., 2018). This section explores the underlying mechanisms of these issues to motivate the development of our ceTDP approach.

The first issue is associated with the use of an overly lenient cluster-defining threshold (CDT) (e.g., *z ≥* 2.33 or *p ≤* 0.01), which can substantially inflate false positive rates in cluster-based neuroimaging analyses (Eklund et al., 2016). The second problem, typically known as the spatial specificity paradox

(Rosenblatt et al., 2018), is that a significant cluster identified by RFT only ensures that there is at least one truly active voxel, but the full extent and location of activation remain unclear. This often leads to faulty interpretations, e.g., that all voxels within the significant cluster are activated. Finally, circular analysis (or double-dipping) occurs when the CDT is chosen after seeing the data. In the circular practice, the same data is used twice, resulting in post-hoc thresholding that biases the inference.

Notably, Eklund et al. (2019) estimated that over 10% of published fMRI studies rely on these problem-atic approaches, indicating a broad and systemic impact in the neuroimaging literature. This creates an urgent need for accessible, statistically rigorous alternatives that can provide valid inference to overcome these specific limitations.

### 2.2 Proposed Solution: Closed Testing

The closed testing is a powerful statistical framework that guarantees strong control over the FWER. The procedure tests all intersections of hypotheses within a hierarchical (or closed) structure, ensuring valid and admissible FWER control for multiple comparisons (Goeman and Solari, 2011). In the neuroimaging context, closed testing allows inference on every possible subset of voxels within the brain. The key requirement of the procedure is that to reject the null hypothesis for a particular cluster, one must reject the null hypotheses for all larger regions containing that cluster (Marcus et al., 1976; Goeman et al., 2021).

Given that the assumption of exchangeability is plausible, several permutation-based variants of closed testing (Blain et al., 2022; Andreella et al., 2023; Vesely et al., 2023), as well as a parametric alternative (Rosenblatt et al., 2018), have been proposed to mitigate the issues associated with standard cluster inference. However, these approaches tend to yield results that are inconsistent with conventional cluster-extent inference.

Our ceTDP approach (Goeman et al., 2023) is an alternative closed testing method that achieves consistency with conventional cluster-extent inference by constructing a closed testing procedure based on classical RFT-based or permutation-based cluster-extent inference. It does this by integrating the cluster-extent inference result as a “local test” within the closed testing framework. When building the framework based on permutation-based cluster-extent inference, this results in consistency with the permutation-based approach, rather than the conventional one. Although cluster-extent inference is used to test both individual and intersection hypotheses, the overall closed testing framework coordinates these tests to ensure a statistically grounded error control. Both the RFT-based and permutation-based cluster-extent inference methods require an a priori choice of a cluster-defining threshold. The ceTDP method inherits this property and also requires such an a priori chosen threshold, unlike some other closed testing methods (Andreella et al., 2023; Blain et al., 2022; Rosenblatt et al., 2018; Vesely et al., 2023).

This effectively extends the traditional cluster inference approach by embedding it within the closed testing framework, which guarantees that the resulting inferences are either uniformly improved or remain equivalent to the original (Goeman et al., 2021). Consequently, when RFT-based cluster-extent inference is used as the local test, ceTDP preserves the conclusions of conventional cluster-level inference while providing additional quantitative information (Goeman et al., 2023).

### 2.3 Beyond Inference: Activation Quantification via TDP

A major advantage of the closed testing framework is that it additionally provides simultaneous lower confidence bounds on the TDP for arbitrary brain regions. For any given non-empty region *V*, ceTDP provides a 100(1 − *α*)% lower confidence bound for its TDP, where the TDP represents the proportion of truly active voxels contained within *V* . Specifically, the TDP lower bound, denoted by *π*(*V*), is defined as

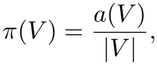

where |*V* | is the total number of voxels in *V*, and *a*(*V*) represents the confidence lower bound on the true discovery number (TDN), i.e., the number of truly active voxels contained within the region *V* . Thus, at *α* = 0.05, a TDP lower bound of *π*(*V*) indicates that at least 100 *π*(*V*)% of the voxels in *V* are truly active with 95% confidence. More importantly, these bounds are valid simultaneously over all possible clusters (or brain subsets). This simultaneity ensures strict control of the cluster-level FWER at level *α*, and thus allows valid, unrestricted post hoc exploration across clusters.

For example, a TDP lower bound of 0.7 for a cluster at *α* = 0.05 indicates that we can state with 95% confidence that at least 70% of its voxels reflect true activation while controlling the cluster-level FWER at 0.05. This outcome is more informative than a binary *p*-value, as it provides a quantitative characterization of the proportion of true activation within any brain region.

TDP lower bounds align with cluster-wise significance in the following way. Clusters that are cluster-wise significant always have TDP greater than zero, and clusters with TDP greater than zero are always cluster-wise significant.

### 2.4 Efficient Two-Stage Framework: From Theory to Practice

In general, a full implementation of closed testing is computationally infeasible for brain-sized data. To address this challenge, ceTDP employs an efficient two-stage framework that makes computation of the TDP lower bound practical (Goeman et al., 2023).

In Stage 1, a cluster-extent threshold is determined using RFT-based or permutation-based cluster-level inference, given a pre-specified CDT. This threshold serves as the key input for subsequent TDP computation. Based on this threshold and, optionally, anatomical or user-chosen regions, the candidate regions are defined and used in Stage 2.

In Stage 2, the TDP lower bound is computed within candidate regions. Two complementary procedures are available: ceTDP-cons, which provides conservative lower bounds, and ceTDP-heur, which employs a heuristic numerical optimization to obtain an estimate of the TDP lower bound. Unlike ceTDP-cons, ceTDP-heur does not provide the same statistical guarantees, because the heuristic optimization is not guaranteed to find the best solution. Consequently, the resulting TDP estimate may overstate the TDP lower bound when insufficient computational resources are used.

The two-stage framework is first illustrated through a motivating example in Section 3. Its computational details are presented in Section 4, followed by practical implementation procedures in Section 5.

## 3 A Motivating Example

This section presents a complete analysis of a real fMRI dataset. It illustrates the limitations of standard cluster-extent inference and demonstrates the transformative results enabled by the ceTDP framework. We analyze data from a sound discrimination task (Pernet et al., 2015) to compare the two approaches.

### 3.1 Data and Preprocessing

We used data from an auditory processing study involving 218 healthy participants who were required to passively listen to vocal and non-vocal sounds (Pernet et al., 2015). For our demonstration, a sub-set of 30 participants was randomly extracted from the full cohort. Individual functional images were preprocessed using SPM12, and the detailed preprocessing pipeline is described in Pernet et al. (2015).

Individual vocal *>* non-vocal contrast images obtained from first-level analysis were analyzed using a one-sample *t*-test at group level to generate the second-level statistical *t*-map and *p*-map for both standard and ceTDP analyses.

### 3.2 Limitation: Low Spatial Specificity of Standard Inference

We first applied standard RFT-based cluster-extent inference (using a CDT of *p ≤* 0.001, and FWER correction at *α* = 0.05). As shown in the first eight columns of Table 1, three clusters were identified significant.

**Table 1:** Cluster-extent inference results for the auditory fMRI vocal *>* non-vocal task contrast using standard RFT-based cluster-extent inference and ceTDP. The table reports all significant clusters, their characteristics, and the TDP bounds obtained using ceTDP-cons and the corresponding TDP estimates obtained using ceTDP-heur. Clusters are listed in descending order of size.

| Cluster |  |  |  | Peak<sup>*</sup> |  |  |  | TDP (%) |  |
|---|---|---|---|---|---|---|---|---|---|
| id | size | brain region | *p*-value | *x* | *y* | *z* | *Z*<sub>max</sub> | ceTDP-cons | ceTDP-heur |
| 1 | 6422 | Right HG/STG/IFG/PT | < 0.001 | 60 | −12 | 2 | 7.22 | 20.5 | 29.6 |
| 2 | 4655 | Left HG/STG/PT | < 0.001 | −60 | −24 | 4 | 6.90 | 22.3 | 31.2 |
| 3 | 329 | Left IFG | < 0.001 | −58 | 16 | 28 | 4.07 | 1.8 | 15.8 |

<sup>\*</sup>Peak MNI coordinates and maximum *z*-score are listed for each cluster.

While this result confirms the presence of activation, it produces a binary map. Each cluster is declared uniformly significant, which raises a critical unresolved question: how much of each cluster is truly active? The lack of spatial specificity is the fundamental limitation that motivates the development of ceTDP.

### 3.3 Solution: Applying the ceTDP Framework

To address the lack of spatial specificity inherent in standard cluster-level inference, we applied the proposed two-stage ceTDP framework to the same statistical maps using the identical CDT of *p ≤* 0.001. Consistent with the conventional analysis, the procedure reproduced the critical cluster-extent threshold of *k* = 129 voxels at *α* = 0.05.

Additionally, this framework provides two complementary measures of the proportion of true activation within significant clusters, namely the conservative TDP lower bound (ceTDP-cons) and a heuristic estimate of the TDP lower bound (ceTDP-heur). These quantities extend binary declaration of cluster significance by quantifying the amount of true activation contained within each significant cluster. The conservative approach provides a hard mathematical guarantee at the cost of being conservative, while the heuristic approach uses numerical optimization and may sometimes overstate the TDP.

### 3.4 Results: Quantitative True Activation Profiles vs Binary Maps

The output of ceTDP fundamentally changes the interpretation of the significant clusters identified by standard cluster-level inference. Accordingly, Table 1 is augmented with two additional columns for each cluster, reporting the conservative TDP lower bound (ceTDP-cons) and the heuristic TDP estimate (ceTDP-heur).

For instance, Cluster 1 (the largest cluster) in Table 1 exhibits a conservative TDP lower bound of *π*_1_ = 20.5% (ceTDP-cons), indicating that, with 95% confidence, at least 20.5% of its voxels correspond to true activation. The corresponding ceTDP-heur estimate is *π*_2_ = 29.6%, providing a heuristic estimate of the TDP lower bound for the same cluster. These relatively low TDP values imply that a substantial portion of the cluster may not represent true activation, illustrating how conventional binary inference can create an overly optimistic impression of spatial specificity.

These results highlight the key contribution of ceTDP. Under conventional cluster-level inference, Cluster 1 would simply be declared significant. In contrast, ceTDP reveals that true activation cannot be claimed to be widely prevalent throughout the cluster. Thus, rather than providing a non-specific binary cluster map, the method yields a quantitative true activation profile, offering substantially improved spatial specificity.

### 3.5 Visualization: From Binary Maps to Quantitative Cluster Maps

The cluster-wise TDP measures can be visualized directly on the brain atlas to complement the binary significance maps produced by conventional inference. In Figure 1, each significant cluster is colorcoded according to its associated TDP lower-bound value, providing an intuitive representation of the estimated minimum proportion of truly active voxels within that cluster.

**Figure 1.**
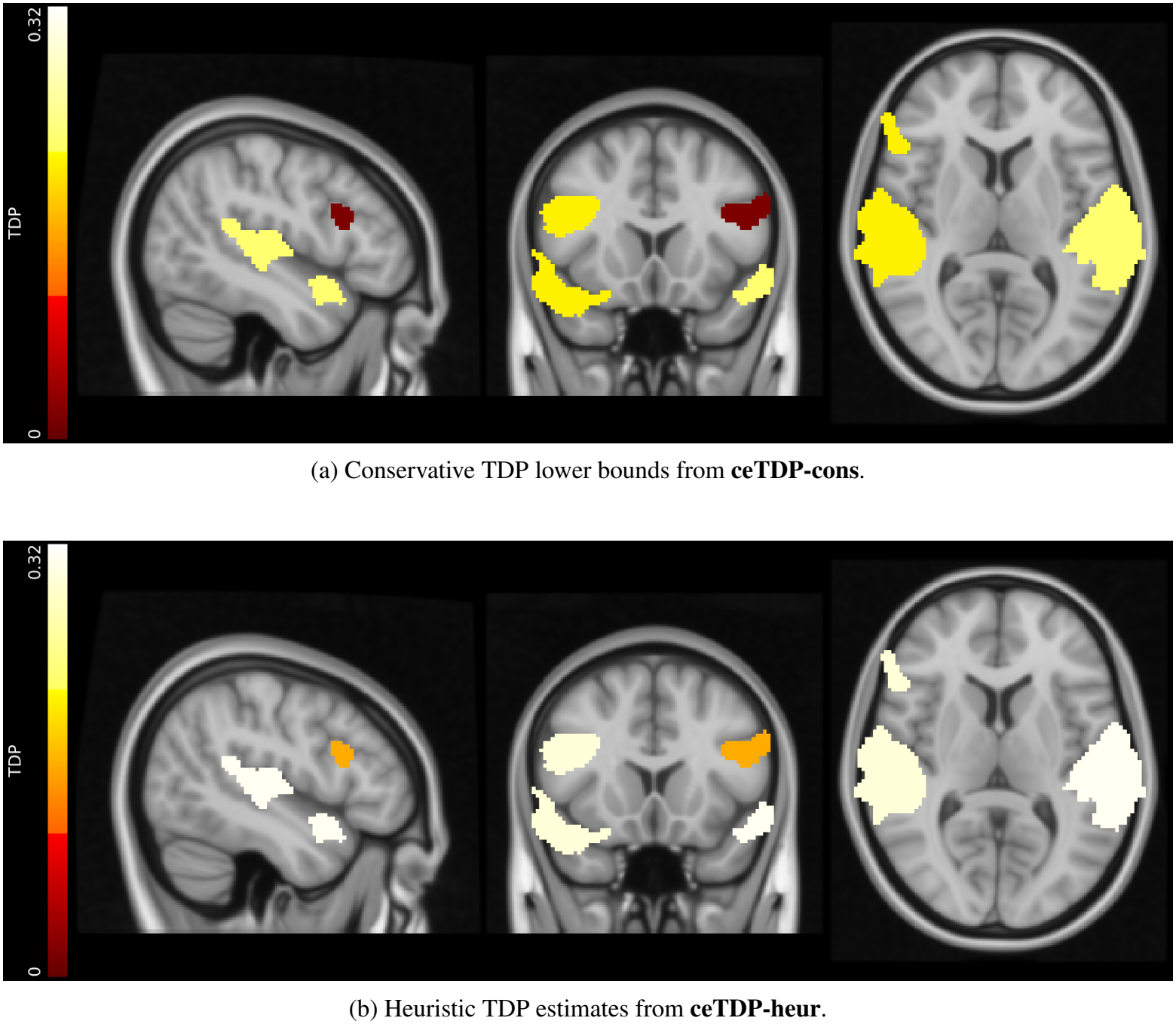
Quantitative cluster maps for the vocal *>* non-vocal contrast used in the motivating example. The three significant clusters are color-coded according to their cluster-wise TDP measures. Separate maps show the results from ceTDP-cons and ceTDP-heur, illustrating the complementary information provided by the two procedures. Clusters are overlaid on the MNI anatomical brain atlas.

Separate visualizations are provided for the conservative TDP lower bound (ceTDP-cons) and the heuristic estimate (ceTDP-heur), illustrating the complementary information offered by the two procedures. A more comprehensive comparison integrating both TDP variants is presented in Figure 2 (see Section 6.1.5).

**Figure 2.**
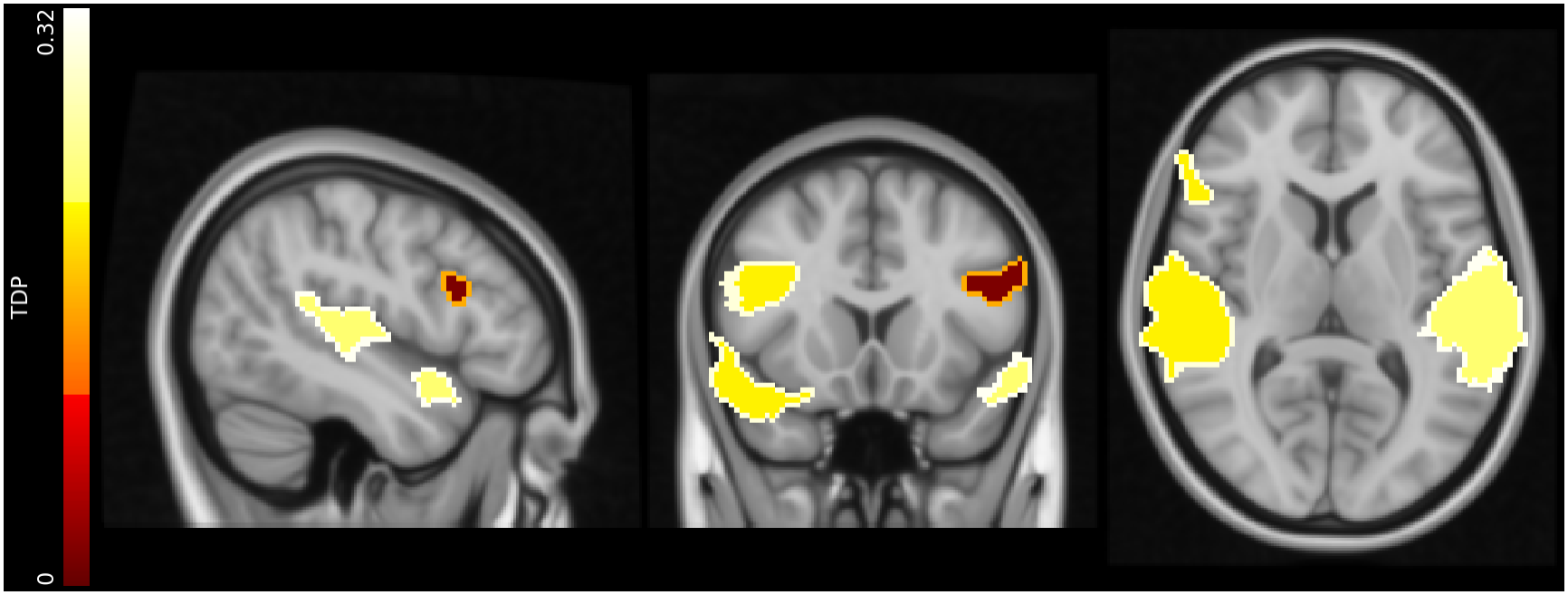
Dual-coding visualization of complementary TDP measures. Clusters are overlaid on an anatomical reference (e.g., the MNI brain atlas). Fill intensity encodes the conservative TDP lower bound (ceTDP-cons), while the cluster outline encodes the heuristic TDP estimate (ceTDP-heur).

### 3.6 From Demonstration to Implementation

This motivating example illustrates how ceTDP extends conventional binary cluster inference by quantifying the proportion of truly active voxels within significant clusters. Having illustrated the additional information provided by this approach, we now turn to its computational implementation.

The next section details the two-stage framework and the complementary conservative (ceTDP-cons) and heuristic (ceTDP-heur) procedures that underlie the results presented above. Their computational and statistical characteristics are examined subsequently.

## 4 The Two-Stage ceTDP Framework

Having established the additional information offered by ceTDP through the motivating example, we now examine the computational framework underlying the approach. The method consists of two sequential stages. Stage 1 determines the cluster-extent threshold inherited from standard cluster-level inference, while Stage 2 obtains the TDP lower bound within candidate regions.

Within Stage 2, as mentioned before, two complementary procedures are available. The conservative procedure, ceTDP-cons, provides theoretically guaranteed TDP lower bounds, whereas the heuristic procedure, ceTDP-heur, employs numerical optimization to obtain potentially tighter estimates of the TDP lower bound.

### 4.1 Stage 1: Determining the Cluster-Extent Threshold

Stage 1 corresponds to the conventional cluster-level inference procedure used in standard neuroimaging analyses. Starting from a chosen CDT, a whole-brain null distribution is obtained using either parametric RFT or permutation-based nonparametric procedures. At a pre-specified significance level *α*, this stage yields the cluster-extent threshold *k*, representing the minimum cluster size required for significance.

Importantly, ceTDP does not replace conventional cluster inference. Instead, it builds directly upon the threshold *k* established at this stage, which subsequently serves as the key input for Stage 2.

#### 4.1.1 Defining Candidate Regions

TDP estimation may subsequently be performed within supra-threshold clusters (STCs), anatomical regions, or arbitrary user-defined regions of interest (ROIs). For each candidate region, its overlap with the significant STCs identified during standard cluster inference is determined. These overlapping regions define the spatial domains within which subsequent TDP inference is carried out.

#### 4.1.2 Preparing Connected Components

Within each candidate region, only connected components contained in its overlap with the significant STCs are retained for further analysis. These contiguous voxel sets constitute the effective search regions on which Stage 2 TDP inference is performed.

Candidate regions containing at least one connected component whose size equals or exceeds the cluster-extent threshold *k* may yield non-zero TDP lower bounds. In contrast, regions for which all connected components are smaller than *k* are immediately assigned a TDP lower bound of zero.

Conceptually, Stage 1 therefore transforms the original statistical image into a collection of contiguous voxel sets that serve as the inputs to the Stage 2 procedures. Practical procedures for generating these connected-component representations are described in Section 5.

### 4.2 Stage 2: Estimating TDP Lower Bounds

Once the threshold *k* and candidate regions have been defined, Stage 2 estimates the TDP lower bound. The procedure first computes lower bounds on the true discovery number (TDN) and subsequently normalizes these values by region size to obtain the corresponding TDP lower bounds.

#### 4.2.2 ceTDP-cons: A Guaranteed Baseline

The ceTDP-cons procedure provides a deterministic and non-iterative lower bound on the proportion of truly active voxels within each region. Owing to the simultaneity property of closed testing, the resulting TDP lower bounds offer strong FWER control and are statistically guaranteed (Goeman et al., 2021).

Consequently, with 100(1 − *α*)% confidence, the reported TDP lower bound is a strict and provable lower bound on the true proportion of active voxels.

From a computational perspective, ceTDP-cons is highly efficient and requires no optimization. It therefore provides a rapid baseline for all candidate regions.

However, this statistical robustness is achieved at the expense of sensitivity. Because the procedure accounts for worst-case configurations arising from factors such as the chosen CDT, image smoothness, search volume, and cluster geometry, the resulting bounds can be conservative. In general, ceTDP-cons tends to be less conservative for large and compact regions and increasingly stringent for small or irregularly shaped regions.

#### 4.2.2 ceTDP-heur: Trading Computation for Sensitivity

The ceTDP-heur procedure builds upon the threshold *k* established in Stage 1 and employs heuristic optimization to search for potentially tighter estimates of the TDP lower bound. By allocating additional computational effort, the algorithm can often achieve substantially higher sensitivity than the conservative baseline.

Unlike ceTDP-cons, the optimization is iterative and heuristic. Although the algorithm is designed such that its estimates never fall below the ceTDP-cons baseline, convergence to the global optimum is not guaranteed. Consequently, the quality of the resulting TDP estimate depends on the computational resources allocated to the optimization procedure.

## 5 Practical Implementation

This section describes the practical workflow for applying the ceTDP framework and summarizes the available software implementations supporting each stage of the analysis. The software suite is designed as a modular ecosystem that integrates cluster-extent threshold computation, connected-component extraction, TDN/TDP lower-bound estimation, and visualization across multiple programming environments.

Users may combine implementations across environments while maintaining a consistent analytical workflow. We first summarize the available software tools and their methodological roles, followed by a high-level description of the workflow and software requirements. Finally, an example workflow is provided to illustrate a typical ceTDP analysis reproducing the motivating example presented in Section 3.

### 5.1 Software Ecosystem

The ceTDP framework is implemented through a collection of interoperable tools distributed across MATLAB/SPM, R/Rcpp, Python, C/C++, and FSL. Together, these implementations support all major components of the analytical workflow, including cluster-extent threshold computation, connected-component extraction, TDN/TDP lower-bound estimation, and visualization of ceTDP results.

In addition to the core inferential procedures, dedicated tools are provided for cluster export and visualization, facilitating integration with arbitrary ROI definitions and external neuroimaging workflows.

Table 2 summarizes the software implementations currently available within the ceTDP framework. These modular tools may be combined across implementations while maintaining methodological consistency throughout the workflow.

**Table 2:** Summary of available software implementations for the ceTDP workflow.

| **Component** | **Method** | **Tool** | **Implementation** | **Interface** |
|---|---|---|---|---|
| **Stage 1** | Parametric *k* | spm_ceTDP_minsize.m | MATLAB / SPM12 | GUI / CLI |
|  |  | smoothest & cluster | FSL | CLI |
|  | Nonparametric *k* | kval.R | R / Rcpp | CLI |
|  | Cluster export | cluster_export.py | Python | CLI |
|  |  | cluster_export.m | MATLAB / SPM12 | GUI / CLI |
| **Stage 2** | ceTDP-cons | spm_ceTDP_cons.m | MATLAB / SPM12 | GUI / CLI |
|  |  | lb | C | CLI |
|  |  | lb.R | R | CLI |
|  |  | lb.py | Python | CLI |
|  | ceTDP-heur | fgreedy | C | CLI |
| **Visualization** | Layered visualization | cetdp_vis.py | Python | CLI |

**Note:** CLI, command-line interface; GUI, graphical user interface; Rcpp, an R package for interfacing R with C++ code.

### 5.2 Workflow Components

The ceTDP workflow consists of four major components:

1. Stage 1 cluster-extent threshold computation.
2. Stage 1 connected-component extraction.
3. Stage 2 TDN/TDP lower-bound estimation (ceTDP-cons and/or ceTDP-heur).
4. Visualization of ceTDP results.

#### 5.2.1 Stage 1: Cluster-Extent Threshold Computation

Stage 1 tools compute the cluster-extent threshold *k*, which serves as the primary input for subsequent TDP lower-bound estimation procedures. Both parametric and nonparametric implementations are available.

The MATLAB/SPM implementation computes *k* directly from SPM statistical models using RFT-based cluster inference. An alternative parametric workflow is provided through the FSL utilities `smoothest` and `cluster`. A nonparametric permutation-based implementation is additionally available in R.

All implementations produce a cluster-extent threshold that can subsequently be supplied to Stage 2.

#### 5.2.2 Stage 1: Connected-Component Extraction

Component extraction utilities are provided to obtain connected components from statistical NIfTI images.

The utilities `cluster export.py` and `cluster export.m` support extraction of whole-brain STCs and ROI-overlapping connected components from statistical NIfTI images, and export voxel-coordinate representations suitable for downstream ceTDP analyses.

The resulting connected-component representations are required for Stage 2 and can be derived either directly from significant STCs or from their overlap with specified regions. This facilitates integration of ceTDP with arbitrary region definitions and external neuroimaging workflows.

#### 5.2.3 Stage 2: TDN/TDP Lower-Bound Estimation

Stage 2 implementations estimate lower bounds on the TDN, from which TDP lower bounds are obtained by normalization with respect to region size.

Implementations of the conservative procedure (ceTDP-cons) are available in MATLAB/SPM, C, R, and Python. The heuristic procedure (ceTDP-heur) is implemented as a standalone C tool based on heuristic optimization.

Together, these implementations provide complementary approaches to quantifying the TDP lower bound within candidate regions.

#### 5.2.4 Visualization

The framework additionally includes visualization utilities for generating NIfTI images containing cluster-level TDP lower bounds and heuristic TDP estimates. These outputs can be overlaid on anatomical reference images using standard neuroimaging visualization tools.

These tools facilitate interpretation of ceTDP results and allow direct comparison between conventional binary cluster maps and quantitative TDP-based representations.

Visualization is an optional component of the workflow and is primarily intended to support interpretation and reporting of results.

### 5.3 Software Requirements and Availability

The software requirements depend on the specific implementations selected for each workflow component. Because the framework is modular, users only need to install the environments corresponding to the tools they intend to use.

MATLAB-based implementations require MATLAB and SPM12. The R-based implementations require a standard R installation together with the necessary supporting packages. The Python-based utilities rely on commonly used scientific-computing libraries. The standalone C/C++ implementations require a suitable compilation environment.

Detailed installation instructions, software dependencies, source code, and usage examples are available through the project repository described in Section 7.

### 5.4 Example Workflow

A typical ceTDP analysis follows a four-step pipeline:

1. **Cluster-extent threshold estimation** The cluster-extent threshold *k* is computed using a selected Stage 1 implementation (parametric or nonparametric).
2. **Definition of candidate regions** Candidate regions are specified as STCs, anatomical ROIs, or arbitrary user-defined regions.
3. **Connected-component extraction** Candidate regions are represented as connected components, either directly from significant STCs or from their overlap with specified regions.
4. **TDP estimation and visualization** ceTDP-cons and/or ceTDP-heur are applied to obtain conservative TDP lower bounds and/or heuristic TDP estimates. These results can optionally be visualized on anatomical images using standard neuroimaging visualization tools. This workflow reproduces the motivating example presented in Section 3. The resulting outputs include the cluster-extent threshold *k*, conservative TDP lower bounds from ceTDP-cons and/or heuristic TDP estimates from ceTDP-heur, together with optional visualizations summarizing the proportion of true activation within significant clusters. Further guidance on interpretation, reporting, and practical use of ceTDP-cons and ceTDP-heur is provided in Section 6.

## 6 Best Practices and Common Pitfalls

This section provides guidance on analytical choices and highlights common pitfalls to ensure reliable and interpretable use of the ceTDP framework.

### 6.1 Best Practices

For robust and transparent ceTDP analyses, we recommend the following practices.

#### 6.1.1 Standardized Reporting of TDP

Standard cluster inference results in neuroimaging are often reported only in terms of *p*-values or binary significance, which provide limited information about the proportion of truly active voxels contained within clusters.

To improve interpretability and reproducibility, cluster-level results should always be accompanied by quantitative estimates of the true activation proportion. Specifically, conservative TDP lower bounds (ceTDP-cons) should be reported alongside conventional statistical results, with heuristic TDP estimates (ceTDP-heur) reported whenever available. In addition, the cluster-extent threshold *k* used in the analysis should always be reported, as it directly determines both cluster significance and subsequent TDP estimation. Reporting *k* also greatly helps in for example, re-analysis or meta-analysis, thereby improving reproducibility in neuroimaging studies.

Together, these quantities transform cluster-level inference from a binary decision framework into a quantitative characterization of cluster-level activation.

#### 6.1.2 Selection of CDT

The CDT plays a central role in cluster inference, as it affects the validity of RFT assumptions and the resulting cluster-extent threshold *k*.

Using overly lenient thresholds (e.g., *z ≥* 2.33 or *p ≤* 0.01) may violate RFT assumptions and lead to inflated false positive rates (Eklund et al., 2016). A commonly recommended choice is *z ≥* 3.09 or *p ≤* 0.001, which is widely adopted in SPM and FSL pipelines.

The choice of CDT also indirectly influences cluster size and the resulting TDP estimates. More lenient thresholds tend to produce larger clusters with lower TDP, while more stringent thresholds yield smaller but more spatially specific clusters.

#### 6.1.3 Joint Interpretation of Conservative and Heuristic TDP Measures

The ceTDP framework provides two complementary measures: the conservative TDP lower bound (ceTDP-cons) and the heuristic TDP estimate (ceTDP-heur).

The conservative TDP lower bound provides a statistically guaranteed minimum proportion of truly active voxels, while the heuristic TDP estimate is typically tighter but does not provide the same formal guarantee.

Both measures should be interpreted jointly. The inequality *π*_cons_ *≤ π*_heur_ always holds, reflecting the fact that ceTDP-heur is designed to improve upon the conservative baseline. A large gap may happen, especially for non-compact clusters.

For transparent reporting and more informative interpretation, both TDP measures should be reported together whenever available.

#### 6.1.4 Candidate Region Construction

Candidate regions may be defined as STCs, anatomical regions, or user-defined ROIs.

For ROI-based or user-defined regions, only voxels overlapping with significant STCs should be included in the analysis. These overlapping voxels are then decomposed into connected components, which serve as the effective input units for Stage 2.

Regions whose STC overlap does not contain any connected component of size at least *k* receive a TDP lower bound of zero and are typically excluded from further interpretation.

#### 6.1.5 Visualization and Reporting

For interpretability, both ceTDP measures should be presented together whenever possible. A recommended strategy is dual-coding:

- cluster fill represents the conservative bound (ceTDP-cons),
- cluster outline represents the heuristic estimate (ceTDP-heur).

This representation distinguishes a conservative lower bound from a heuristic estimate of the proportion of truly active voxels.

A standard reporting template should include cluster size, peak coordinates, CDT, cluster-extent thresh-old *k*, and the conservative TDP lower bound and heuristic TDP estimate.

Example reporting template:

Applying a cluster-defining threshold of *p ≤* 0.001 (or *t ≥* 3.40), we identified three significant clusters using RFT-based cluster inference with *k* = 129 at *α* = 0.05. For Cluster 1 (6422 voxels; peak MNI (60, −12, 2)), ceTDP-cons indicated that, with 95% confidence, at least 20.5% of voxels were active, while ceTDP-heur estimated the corresponding TDP lower bound of 29.6%. Similar patterns were observed for Clusters 2 and 3.

These results are visualized in Figure 2 using the dual-coding strategy.

### 6.2 Common Pitfalls and Recommended Solutions

#### 6.2.1 Misinterpreting Binary Cluster Inference

Cluster-level inference only guarantees that at least one voxel within a cluster is active. It does not imply that all voxels are active.

Interpreting clusters as homogeneous leads to overestimation of spatial specificity and overstatement of inferential precision. ceTDP addresses this limitation by quantifying the proportion of truly active voxels within each cluster.

#### 6.2.2 Misuse of Candidate Regions and STC Overlap

Valid TDP estimation requires restricting analysis to voxels that overlap with significant STCs. Including non-overlapping voxels can lead to invalid or inflated estimates.

For ROI-based analyses, candidate regions must first be intersected with STCs, and only resulting connected components should be used for inference. Regions without connected components of size at least *k* are assigned a lower bound of zero for the TDP.

#### 6.2.3 Misinterpreting ceTDP-heur as a Guaranteed TDP Lower Bound

The ceTDP-heur procedure is based on heuristic optimization and does not guarantee convergence to the global optimum. Although it is constrained to never fall below ceTDP-cons, it should not be interpreted as a strictly valid inferential lower bound on its own.

Reliable interpretation requires joint consideration of both ceTDP-cons and ceTDP-heur.

#### 6.2.4 Inadequate Runtime for ceTDP-heur

Because ceTDP-heur relies on iterative optimization, insufficient runtime may lead to suboptimal heuristic estimates.

Empirical evaluations suggest that stable convergence is typically achieved with sufficiently relaxed stopping criteria and extended runtime. A recommended configuration is:
   

~~~
./fgreedy -x batch7.cnf -T0 -N60 -k*<*value*> <*file*>*
~~~

where -T0 allows unlimited runtime and -N60 specifies a 60-second no-improvement threshold per iteration.

Under these settings, ceTDP-heur typically converges to stable estimates for most practical datasets.

## 7 Conclusion and Discussion

This tutorial paper introduces ceTDP, a two-stage framework that extends conventional RFT-based (parametric) and permutation-based (nonparametric) cluster inference. ceTDP addresses key limitations of standard cluster-level inference, including limited spatial specificity and the risk of circular analysis.

By integrating cluster inference with closed testing, the framework provides strong control of the FWER while additionally enabling quantification of activation via the TDP. Its two-stage design combines a statistically rigorous procedure (ceTDP-cons) with a heuristic optimization procedure (ceTDP-heur), providing complementary measures of true activation. Importantly, both variants remain consistent with standard cluster inference when used as a local test.

A key contribution of ceTDP is that it complements binary significance statements with quantitative estimates of the proportion of truly active voxels within clusters. This shifts the interpretation of cluster-level results from a binary decision framework to a quantitative characterization of cluster-level activation. Such information enables more nuanced interpretation of cluster findings and facilitates more consistent reporting across studies.

In summary, ceTDP provides a practical framework for augmenting conventional cluster inference with statistically grounded measures of true activation. Through its modular software implementations, the method is accessible within standard neuroimaging workflows. We expect that its use can improve the interpretability and quantitative reporting of cluster-level results in neuroimaging.

## Data and Code Availability

The implementation of the ceTDP framework, along with the example dataset described in Section 3, is publicly available on GitHub. The ceTDP project is organized around a main GitHub repository (https://github.com/xuchen312/ceTDP), with individual tools maintained in their respective repositories. The example fMRI dataset used in this paper was obtained from the OpenNeuro repository and is publicly available under accession number ds000158 (Pernet et al., 2015).

## Acknowledgements

This work was supported by the London Mathematical Society Scheme 4: Research in Pairs grant (Ref. 42438 to X.C.).

